# Two central pattern generators from the crab, Cancer borealis, respond robustly and differentially to extreme extracellular pH

**DOI:** 10.1101/374405

**Authors:** Jessica A. Haley, David Hampton, Eve Marder

**Affiliations:** Volen Center and Biology Department, Brandeis University, Waltham, MA 02454

**Keywords:** stomatogastric ganglion, cardiac ganglion, pyloric rhythm, crustacean, ocean acidification

## Abstract

Animals and their neuronal circuits must maintain function despite significant environmental fluctuations. The crab, *Cancer borealis*, experiences daily changes in ocean temperature and pH. Here, we describe the effects of extreme changes in extracellular pH – from pH 5.5 to 10.4 – on two central pattern generating networks, the stomatogastric and cardiac ganglia of *C. borealis*. Given that the physiological properties of ion channels are known to be sensitive to pH within the range tested, it is surprising that these rhythms generally remained robust from pH 6.1 to pH 8.8. Unexpectedly, the stomatogastric ganglion was more sensitive to acid while the cardiac ganglion was more sensitive to base. Considerable animal-to-animal variability was likely a consequence of similar network performance arising from variable sets of underlying conductances. Together, these results illustrate the potential difficulty in generalizing the effects of environmental perturbation across circuits, even within the same animal.

**Abbreviations:**

STG: stomatogastric ganglion
CG: cardiac ganglion
CPG: central pattern generator
AB: Anterior Burster
PD: Pyloric Dilator
LP: Lateral Pyloric
PY: Pyloric
SC: Small Cell
LC: Large Cell
*lvn*: lateral ventricular nerve
ANOVA: analysis of variance
PTX: picrotoxin
IPSP: inhibitory post-synaptic potential
LG: Lateral Gastric
MG: Medial Gastric
LPG: Lateral Posterior Gastric
GM: Gastric Mill
DG: Dorsal Gastric
AM: Anterior Median
Int1: Interneuron 1
*mvn*: medial ventricular nerve
*dgn*: dorsal gastric nerve
*lgn*: lateral gastric nerve
*ion*: inferior oesophageal nerve
IC: Inferior Cardiac
VD: Ventricular Dilator
MCN1: Modulatory Commissural Neuron 1
VCN: Ventral Cardiac Neuron
CPN2: Commissural Projection Neuron 2
CoG: commissural ganglion
KDE: kernel density estimate
IQR: interquartile range
CI: confidence interval

## Introduction

Nervous systems must be both robust and adaptable to changes in internal and external conditions. Many intertidal marine crustaceans, such as the crabs and lobsters inhabiting the North Atlantic, experience large fluctuations in ocean temperature, acidity, dissolved oxygen levels, and salinity. The Jonah crab, *Cancer borealis*, can often be found foraging for food in intertidal zones where it experiences temperatures between 3°C and 24°C with fluctuations as great as 10°C in a single day (Donahue et al., 2009; Haeffner Jr, 1977; Stehlik et al., 1991). As pH is temperature-dependent, ocean pH fluctuations occur over the daily, monthly, and yearly experiences of long-lived crustaceans, such as *C. borealis.*

Increased atmospheric carbon dioxide is causing rises in both the temperature (Buchel et al., 1999; Macfarling et al., 2006) and dissolved carbon dioxide concentration of the world’s oceans (Canadell et al., 2007; Knorr, 2009). If current trends continue, estimates reveal that these changes will reduce the average ocean pH from 8.1 to 7.6 in the next century (Wittman and Pörtner, 2013). The effects of ocean acidification are noticeably disrupting marine ecosystems (Kelly and Hofmann, 2013; Webb et al., 2016). In the lobster, *Homarus americanus*, prolonged thermal stress results in pH acidosis, hyperchloremia, and hyperproteinemia (Dove et al., 2005) and has been linked to mortality (Pearce and Balcom, 2005). Further, shifts in hemolymph pH *in vitro* alter the frequency and strength of the lobster cardiac rhythm (Qadri et al., 2007).

In several marine invertebrates including *H. americanus* and the crab, *Carcinus maenas*, hemolymph pH varies inversely with temperature following the rules of constant relative alkalinity (Dove et al., 2005; Howell et al., 1973; Qadri et al., 2007; Truchot, 1973, 1986; Vogt and Regehr, 2001). In other words, as temperature increases, hemolymph acidifies by approximately -0.016 pH/°C to maintain a constant ratio of pH to pOH through a process of bicarbonate buffering. Maintenance of this ratio in extra- and intracellular fluids is thought to be important for stabilizing macromolecular structure and function (Reeves, 1972, 1977; Somero, 1981; Truchot, 2003). Like hemolymph, intracellular pH generally decreases as temperature rises, but has been shown to change at varying rates in different tissues in the crab, *Callinectes sapidus* (Vogt and Regehr, 2001). Active mechanisms for the maintenance of intracellular pH have been suggested in the crab, *Cancer pagurus* (Golowasch and Deitmer, 1993).

Both temperature and pH alter the biophysical parameters governing the activity of ion channels and pumps in excitable membranes. Studies on the biophysical effects of pH on ion channels revealed attenuation of sodium, calcium, and potassium currents under acidic extracellular conditions (Doering and McRory, 2007; Hille, 1968; Mozhaev et al., 1970; Tombaugh and Somjen, 1996; Wanke et al., 1979; Zhou and Jones, 1996). In the frog myelinated nerve, changing extracellular saline from pH 7 to pH 5 reduces sodium currents by up to 60% in a membrane potential-dependent manner (Woodhull, 1973) which may be mediated by increased sodium channel inactivation (Courtney, 1979). In rat CA1 neurons, moderate pH shifts from 7.4 to 6.4 reversibly depressed and shifted the voltage dependence of sodium (15% attenuation; +3 mV shift) and calcium currents (60% attenuation; +8 mV shift). Further, this acidosis was sufficient to shift potassium current inactivation to more depolarized voltages (+10 mV) while moderate alkalosis up to pH 8.0 had the opposite effect (Tombaugh and Somjen, 1996). This reversible gating of sodium, calcium, and potassium channels may result from protonation of an acidic group with pK_a_ between 5.2 and 7.1 (Tombaugh and Somjen, 1996). Decreases in glutamatergic synaptic function (Billups and Atwell, 1996; Bloch et al., 2001; Sinning et al., 2011) and increases in GABAergic synaptic function (Sinning and Hübner, 2013) in response to decreases in pH have also been shown.

Although the biophysical, ethological, and environmental implications of changing pH have been well studied, less is known about the effect of pH changes on neuronal circuits in marine invertebrates. Here, we study the effects of acute changes in extracellular pH on two well characterized neuronal circuits, the stomatogastric (STG) and cardiac (CG) ganglia, of the crab, *C. borealis*. These central pattern generators (CPGs) drive the coordinated and rhythmic muscle movements of the crab’s stomach and heart, respectively. Although these CPGs are driven by numerically small neuronal circuits, their dynamics involve complex interactions between many intrinsic and synaptic currents.

Previous studies have shown that the pyloric rhythm of the STG and cardiac rhythm of the CG are remarkably robust to short- and long-term changes in temperature in both *in vivo* and *ex vivo* preparations (Kushinsky et al., 2018; Marder et al., 2015; Soofi et al., 2014; Tang et al., 2010; Tang et al., 2012). Despite similarly robust activity under moderate perturbation, these experiments have revealed animal-to-animal variability in network activity at extreme temperatures (Haddad and Marder, 2018; Tang et al., 2010; Tang et al., 2012). In accordance with these findings, we reveal that both the pyloric and cardiac rhythms are remarkably robust to acute pH changes from 5.5 to 10.4. This is surprising given the sensitivity of many ion channels to pH in these ranges and suggests that networks can be more robust to pH changes than expected.

## Results

Two neuronal networks were studied in this paper. The stomatogastric ganglion (STG) of the crab, *Cancer borealis*, contains the neurons that generate two stomach rhythms, the fast pyloric rhythm and the slower gastric mill rhythm. The pyloric rhythm is driven by a three-neuron pacemaker kernel – one Anterior Burster (AB) and two Pyloric Dilator (PD) neurons. The Lateral Pyloric (LP) and Pyloric (PY) neurons fire out of phase with the PD neurons because they are rhythmically inhibited by the PD/AB group (Marder and Bucher, 2007). The cardiac ganglion (CG) generates the rhythm responsible for heart contraction, and consists of four pacemaker Small Cell (SC) neurons that drive five motor Large Cell (LC) neurons (Cooke, 2002).

A schematic diagram of the stomatogastric nervous system preparation is found in Figure 1A. Intracellular recordings were made from the somata of the desheathed STG and examples of the LP and PD neuron waveforms are shown. Extracellular recordings from the motor nerves are indicated by the gray circles. An example of the triphasic activity of the LP, PD, and PY neurons is seen in the inset trace next to the lateral ventricular nerve (*lvn*). The connectivity diagram of the major neuron classes of the triphasic pyloric rhythm is given in Figure 1B.

**Figure 1.**
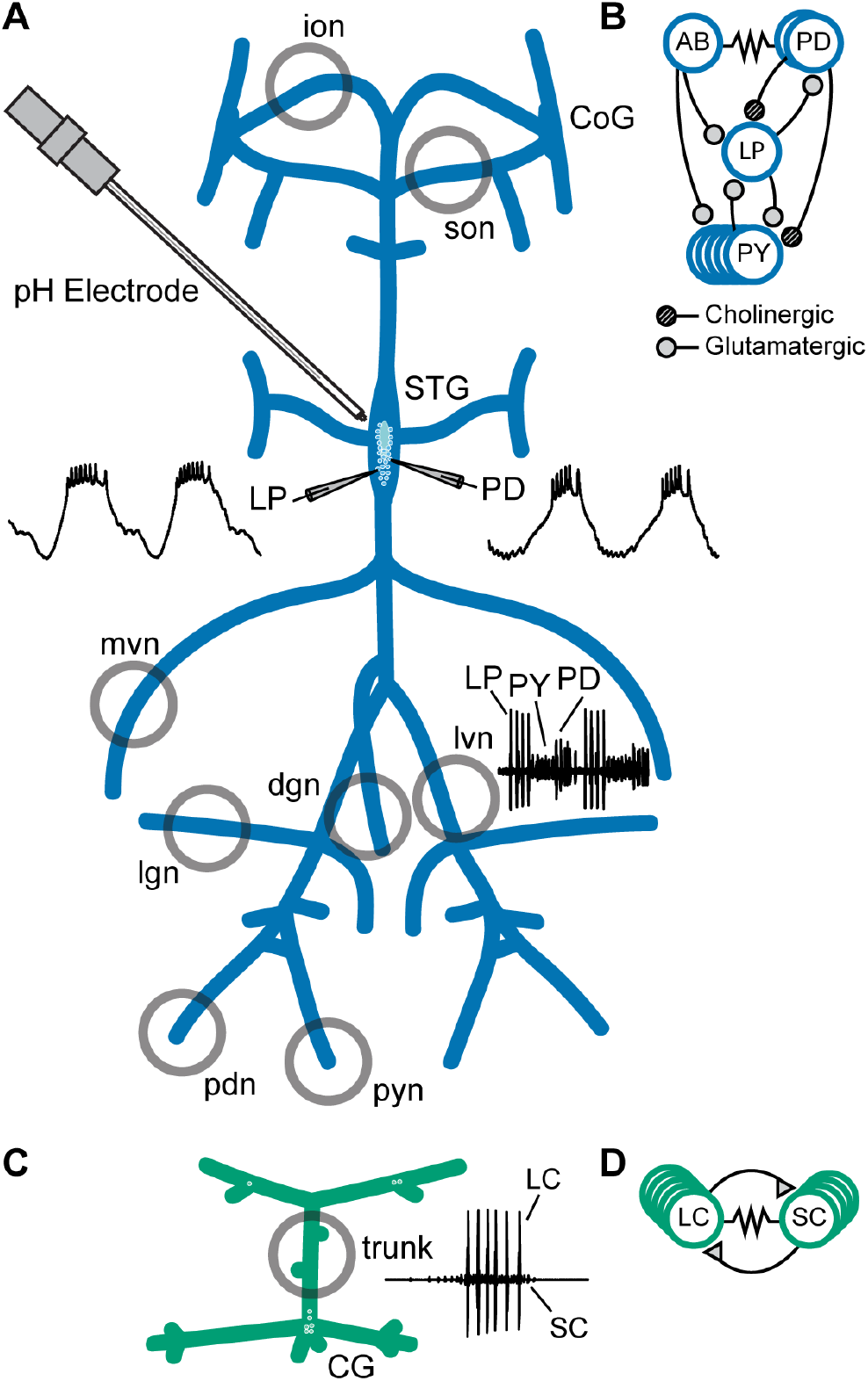
Preparations and circuit diagrams. **(A)** Schematic of the stomatogastric nervous system preparation. Extracellular electrodes were placed in vaseline wells (gray circles) drawn around nerves of interest. An example extracellular nerve recording from the lateral ventricular nerve (*lvn*) shows two cycles of the triphasic pyloric rhythm containing spikes from the Lateral Pyloric (LP), Pyloric (PY), and Pyloric Dilator (PD) neurons. Example intracellular recordings from the LP and PD neurons are displayed. **(B)** Simplified diagram of the pyloric circuit. Filled circles represent inhibitory chemical synapses; resistor symbol represents electrical coupling. **(C)** Schematic of the cardiac ganglion preparation. Extracellular electrodes were placed in a well (gray circle) around the trunk of the preparation. An example extracellular recording shows one burst of the Small Cell (SC) and Large Cell (LC) neurons. **(D)** Diagram of the cardiac circuit. Filled triangles represent excitatory chemical synapses; the resistor symbol represents electrical coupling.

A schematic diagram of the crab cardiac ganglion preparation shows an example of one burst of SC and LC activity recorded from the trunk (Figure 1C). Figure 1D shows the connectivity diagram of the cardiac ganglion.

### The Pyloric rhythm is surprisingly robust to extreme changes in pH

To characterize the response of the pyloric rhythm to acute changes in pH, superfused saline was exchanged every 15 minutes in steps of approximately pH 0.5 from a control pH of 7.8 to an extreme pH of either 5.5 or 10.4. Following the first acid or base step protocol, preparations were allowed to recover for a minimum of 30 minutes at control pH until the frequency of the pyloric rhythm approached that of controls. Preparations were then subjected to a step protocol in the opposite direction followed by a second recovery period. Acid-or base-first protocols were counterbalanced.

Recordings and analysis from an example STG experiment with an acid-first protocol are shown in Figure 2. Each box contains simultaneous intracellular recordings of the PD and LP neurons and extracellular recordings of the *lvn* during the last minute at each pH step (Figure 2A). STG #1 demonstrated a normal triphasic rhythm in control saline at pH 7.8 (Figure 2A; top left). As the preparation was subjected to more acidic saline, the rhythm remained triphasic until the most acidic pH, 5.5. Control activity was recovered when the preparation was once again placed in control saline (Figure 2A; top right). The bottom row shows the same preparation in basic conditions where it remained triphasic, although with fewer spikes per burst and lower amplitude slow waves at pH 10.4. Again, the preparation recovered a canonical triphasic rhythm in control saline as seen in the bottom right.

**Figure 2.**
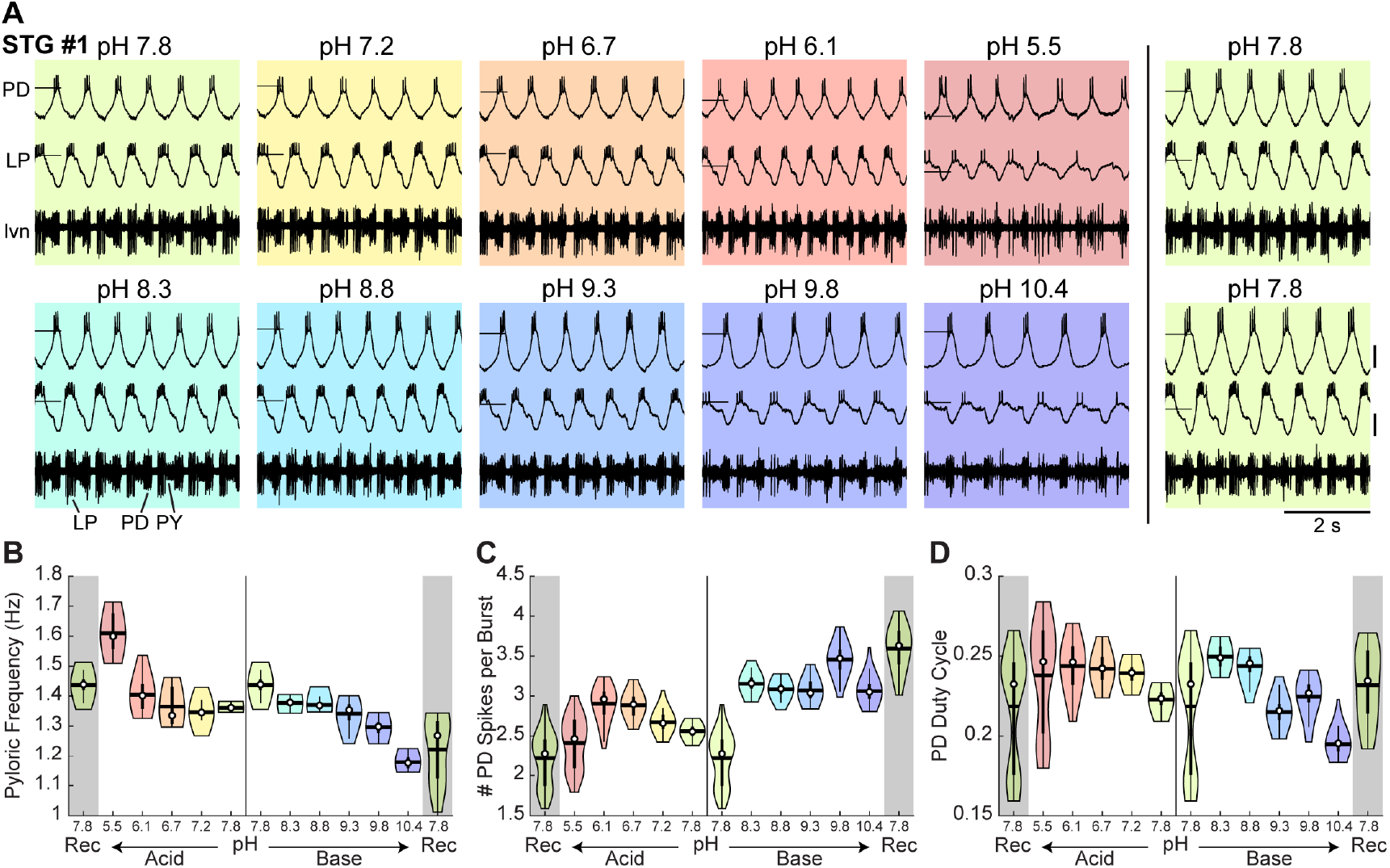
Robust pyloric rhythm activity across pH. **(A)** Example recordings from a stomatogastric ganglion experiment with an acid-first protocol. Intracellular recordings of the PD and LP neurons and extracellular recordings of the *lvn* are shown. Each colored box displays five seconds of recordings taken from the last minute at each pH step. The experiment can be read left to right then top to bottom in chronological order. Horizontal lines indicate a reference membrane potential of -40 mV; vertical lines indicate a scale of 10 mV. **(B)** Pyloric frequency, **(C)** number of PD spikes per burst, and **(D)** PD duty cycle were calculated for the last eight minutes of each pH step. Violin plots show the KDE distribution, mean, median, IQR, and 95% CI for each measure across pH conditions. Recoveries from acid and base are displayed in the shaded gray regions on the far ends of each plot.

Measures of the pyloric rhythm burst frequency, PD spikes per burst, and PD duty cycle (the fraction of the pyloric rhythm’s period during which the PD neurons were active) were calculated for a period of steady state activity (the last eight minutes of each 15-minute pH step). Violin plots reveal the distribution of these measures for STG #1 at each pH (Figure 2B-D). The pyloric burst frequency of STG #1 increased in acid and decreased in base (Figure 2B). The number of PD spikes per burst decreased at pH 5.5 (Figure 2C). The duty cycle of the PD neurons in STG #1 decreased slightly in both acid and base (Figure 2D).

While some preparations maintained surprisingly robust pyloric rhythms from pH 5.5 to 10.4, others exhibited disrupted patterns of activity at the most extreme pH conditions. The activity of two additional STG preparations across the range of pH tested is shown in Figure 3A. Both preparations displayed robust activity across a nearly three-fold range of hydrogen ion concentration but became weakly active or silent at pH 5.5. At pH 10.4, STG #2 was slow and weakly triphasic while STG #3 retained a strong triphasic rhythm. These examples highlight both the animal-to-animal variability of the pyloric rhythm at control conditions and the variable effects of extreme acidosis or alkylosis on this network.

**Figure 3.**
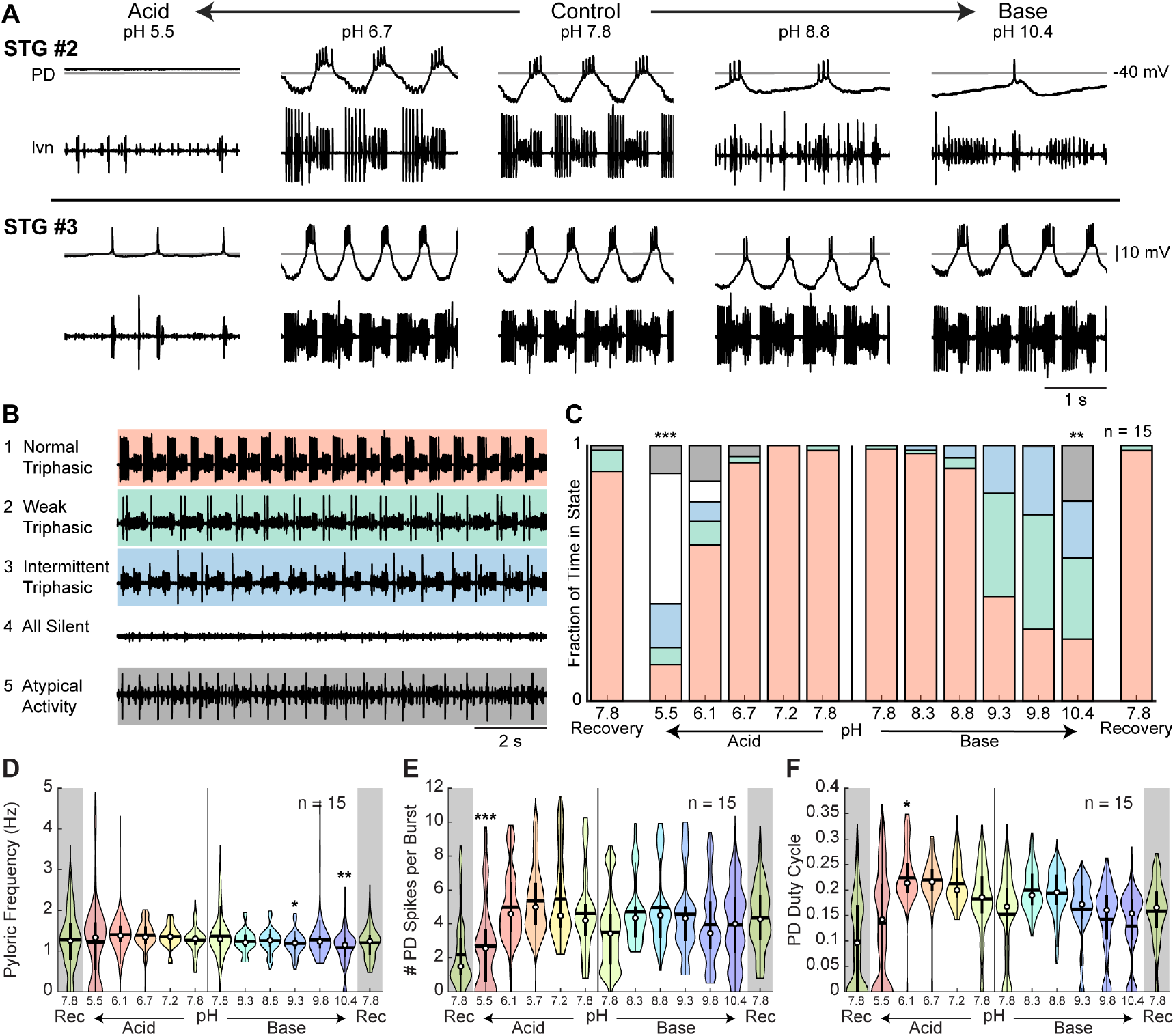
Variability of pyloric rhythm activity at extreme pH. **(A)** Two additional stomatogastric ganglion experiments displaying three seconds of intracellular PD and extracellular *lvn* recordings. Horizontal lines indicate a reference membrane potential of -40 mV; vertical line indicates a scale of 10 mV. **(B)** Five states were defined to characterize pyloric rhythm activity. Examples of activity for each state are given. **(C)** Stacked bars give the mean fraction of time that all 15 preparations spent in each state. **(D)** Pyloric rhythm frequency, **(E)** number of PD spikes per burst, and **(F)** PD duty cycle were calculated and pooled across all STG preparations for each pH step. Violin plots show the KDE distribution, mean, median, IQR, and 95% CI for each measure across pH conditions. Recoveries from acid and base are displayed in the shaded gray regions on the far ends of each plot. Asterisks denote statistical significance revealed by paired samples t-tests with Bonferroni correction (*p<0.05; **p<0.01; ***p<0.001).

To characterize these effects across all preparations, we defined five states of activity: 1) ‘normal triphasic’ rhythm containing PD, LP, and PY with a minimum of three spikes per burst for each unit; 2) ‘weak triphasic’ rhythm retaining all 3 units with some units spiking only once or twice per cycle; 3) ‘intermittent triphasic’ rhythm describing rhythmic activity with only some units active; 4) ‘all silent’; and 5) ‘atypical activity’ or activity that could not be categorized under the first four definitions (Figure 3B). Preparations were categorized systematically according to the criteria outlined in Materials and Methods. The mean fraction of time that all preparations spent in these five states at steady state was analyzed (Figure 3C). Both acid and base significantly decreased the fraction of time that preparations were rhythmic, a combined metric of states 1 (normal triphasic) and 2 (weak triphasic) (Figure 3–figure supplement 1). Preparations were significantly less triphasically rhythmic at pH 5.5 and pH 10.4 compared to control pH 7.8.

To describe these effects quantitatively, measures of the pyloric rhythm frequency, the number of PD spikes per burst, and PD duty cycle were calculated. Violin plots give pooled distributions for each pH across all preparations (Figure 3D-F). Mean pyloric burst frequency was relatively invariant across pH values in the presence of acid, but varied significantly across base steps (Figure 3D). Pyloric burst frequency at pH 9.3 and 10.4 was significantly slower than that at control. Both acid and base significantly affected the mean number of PD spikes per burst (Figure 3E). The number of spikes per burst was significantly reduced at pH 5.5. Additionally, there was a significant effect of acid and base on the mean PD duty cycle (Figure 3F). Paired samples t-tests revealed a slight increase in mean PD duty cycle from control pH 7.8 to pH 6.1.

The pooled distributions for these three measures were highly variable for all pH conditions reflecting the animal-to-animal variability in the pyloric rhythm. We plotted the distributions for all 15 STG preparations at control pH 7.8 and found similarly variable activity at baseline conditions (Figure 3–figure supplement 2).

### Isolated pyloric neurons are sensitive to extreme pH

To determine how the intrinsic properties of isolated neurons respond to pH, we analyzed several characteristics of the intracellular recordings from the PD and LP neurons (Figure 4). We isolated the neurons from most of their pyloric network synaptic inputs by blocking the glutamatergic inhibitory synapses with 10^-5^ M picrotoxin (PTX) (Marder and Eisen, 1984)(Figure 1B). We analyzed mean resting membrane potential (mV), spike amplitude (mV), and burst or spiking frequencies (Hz) for both cells in the presence of PTX as a function of pH. The waveforms of the PD and LP neurons from an example preparation are shown prior to PTX superfusion (Figure 4A; leftmost traces). Note the large LP-evoked inhibitory post-synaptic potentials (IPSPs) in the trough of the PD neuron waveform and the large amplitude inhibition of the LP caused by activity of the PD, AB, and PY neurons. Following application of PTX at control pH 7.8 (Figure 4A; center traces), the PD neuron was still bursting but the LP-evoked IPSPs were entirely blocked. Most of the inhibitory inputs to the LP neuron were blocked, leaving only the cholinergic inhibition contributed by the PD neurons. At pH 6.7, the PD neuron of this preparation lost most of its slow wave activity. It then became silent and depolarized in pH 5.5 saline. At pH 8.8, the PD burst was largely intact, and at pH 10.4, the neuron showed single spike bursts. The LP neuron fired tonically from pH 6.7 to 10.4, again showing loss of activity at pH 5.5 similar to PD.

**Figure 4.**
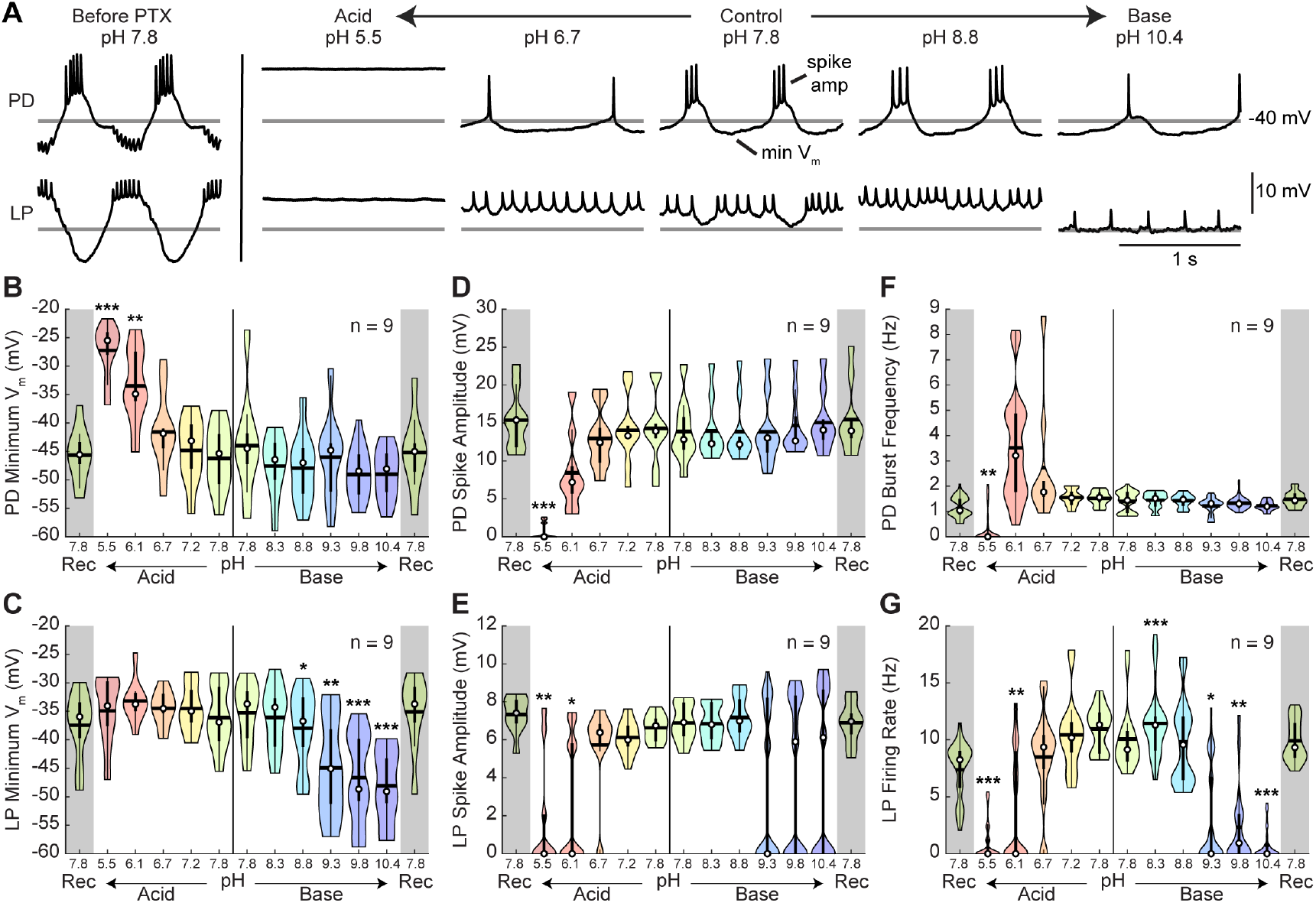
Intracellular characteristics of isolated pyloric neurons, PD and LP. Several characteristics of the PD and LP neurons in the presence of picrotoxin (PTX) were measured for the last minute of each pH condition. **(A)** Example intracellular recordings of PD and LP neurons prior to PTX application and in the presence of PTX across pH conditions. Horizontal lines indicate a reference membrane potential of -40 mV; the vertical line indicates a scale of 10 mV. **(B,C)** Minimum membrane potential and **(D,E)** spike amplitude are plotted for LP and PD as a function of pH. **(F)** PD burst frequency and **(G)** LP firing rate are also plotted at each pH. Violin plots show the KDE distribution, mean, median, IQR, and 95% CI for each measure across pH conditions. Recoveries from acid and base are displayed in the shaded gray regions on the far ends of each plot. Asterisks denote statistical significance revealed by paired samples t-tests with Bonferroni correction (*p<0.05; **p<0.01; ***p<0.001).

Violin plots show pooled values for the most hyperpolarized levels (minimum voltages) of the membrane potential for PD and LP neurons (Figure 4B,C). For moderate shifts in pH, the membrane potential was fairly stable. At extreme acid, the mean PD neuron membrane potential depolarized significantly, while the mean LP neuron membrane potential remained relatively constant (Figure 4–figure supplement 1). In contrast, the PD neuron’s membrane potential in basic saline was relatively constant, even at extreme base, but the LP neuron’s membrane potential significantly hyperpolarized. The depolarization of PD at pH 5.5 and 6.1 and the hyperpolarization of LP at pH 8.8, 9.3, 9.8, and 10.4 were significantly different from control saline.

Additionally, there was a slight effect of pH on mean spike amplitude at the most extreme pH conditions for both the LP and PD neurons (Figure 4D,E). Acid significantly affected both PD and LP neuron spike amplitude while alkylosis had an effect on the LP, but not the PD neurons (Figure 4–figure supplement 1). At pH 5.5 spike amplitude was significantly attenuated for both LP and PD. Additionally, LP spike amplitude was significantly decreased at pH 6.1 while PD was only moderately affected.

There was a significant effect of both acidic and basic saline on mean PD burst frequency and LP firing rate (Figure 4F,G; Figure 4–figure supplement 1). Mean PD burst frequency was significantly decreased at pH 5.5. The LP firing rate was significantly reduced in pH 5.5, 6.1, 8.3, 9.3, 9.8, and 10.4 compared to control pH 7.8.

### Rhythmic gastric-like activity was elicited upon exposure to and recovery from extreme acid and base

The STG contains a second slower central pattern generating circuit known as the gastric mill rhythm (Marder and Bucher, 2007). Unlike the pyloric rhythm which contains a pacemaker kernel, the gastric rhythm is controlled by the reciprocal alternation of activity driven by descending neuromodulatory inputs (Marder and Bucher, 2007; Nusbaum et al., 2017). The principal neurons involved in the gastric mill rhythm are the Lateral Gastric (LG), Medial Gastric (MG), Lateral Posterior Gastric (LPG), Gastric Mill (GM), Dorsal Gastric (DG), and Interneuron 1 (Int1) neurons (Mulloney and Selverston, 1974a, b).

The gastric mill rhythm is often silent in dissected STG preparations and requires stimulation of descending and/or sensory neurons to elicit activity (Blitz et al., 1999). Interestingly, in 10 of 15 preparations a gastric-like rhythm appeared at pH 8.8 or above, and 4 of 15 showed this type of activity in acid at or below pH 6.1. Further, a strong gastric-like rhythm was elicited upon recovery from extreme acid in 5 of 15 preparations and from extreme base in 7 of 15. Preparations in which gastric rhythms were seen in one of these conditions were likely to display gastric-like activity in the other conditions.

An example preparation in control pH 7.8 saline is shown as it recovers activity after exposure to pH 5.5 saline (Figure 5). Intracellular recordings from the LP and PD neurons and extracellular recordings from five nerves – lateral ventricular (*lvn*), medial ventricular (*mvn*), dorsal gastric (*dgn*), lateral gastric (*lgn*), and inferior oesophageal (*ion*) – are shown. In addition to axons of the LP, PD, and PY neurons, the *lvn* contains the LG axon. The *mvn* contains axons from two neurons, the Inferior Cardiac (IC) and the Ventricular Dilator (VD). Inhibition of IC and VD was coincident with LG bursting. The *dgn* shows GM and DG activity and the *lgn* contains LG activity. Recordings from the *ion* reveal Modulatory Commissural Neuron 1 (MCN1) activity. One period of the gastric mill rhythm can be defined by the time from the onset of one LG burst to the next.

**Figure 5.**
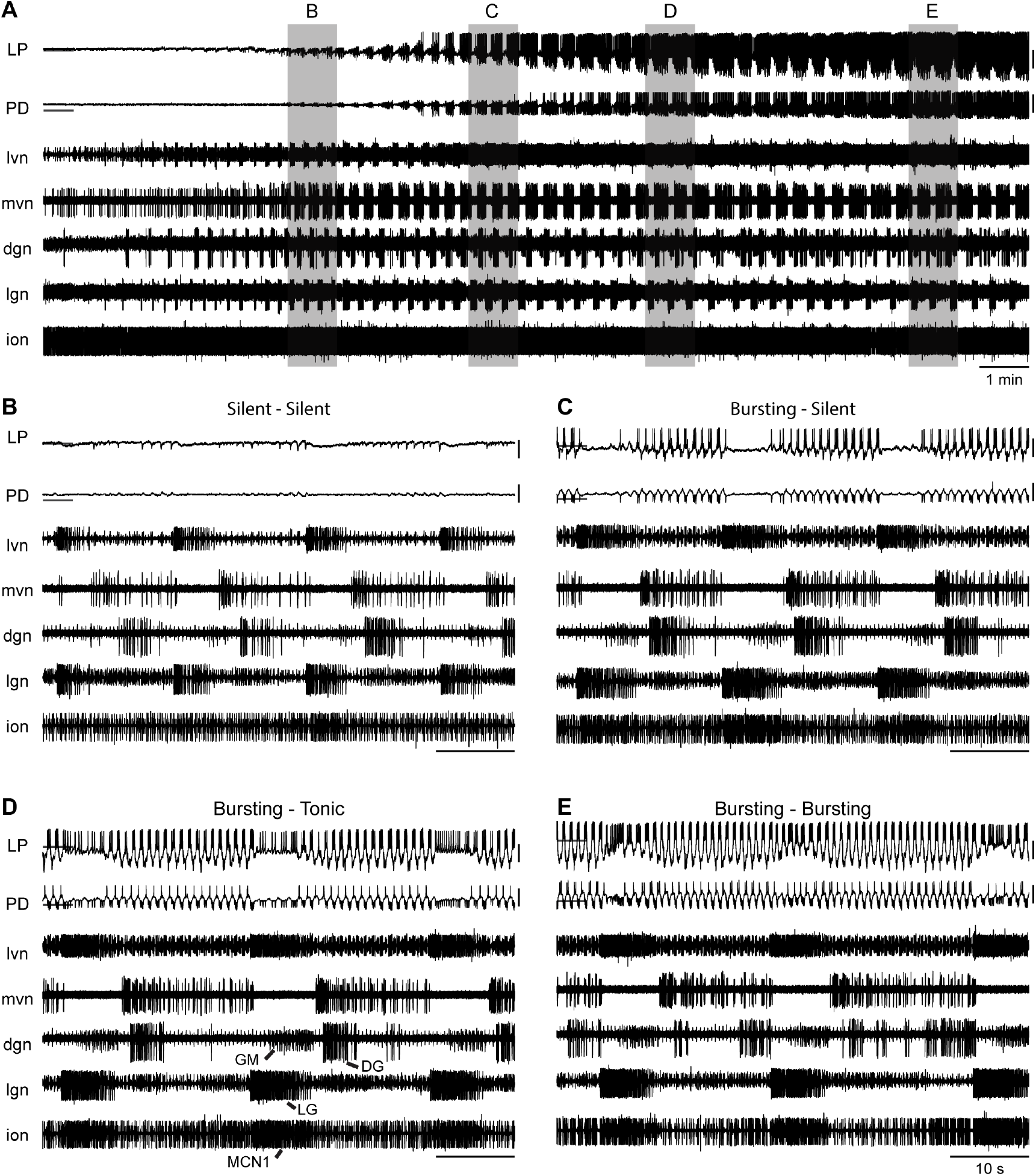
Rhythmic gastric-like activity upon recovery from extreme acid. **(A)** 20 minutes of recording are shown from an example experiment where the ganglion had become silent at pH 5.5 and began recovering rhythmic activity in control pH 7.8 saline. Intracellular recordings from LP and PD neurons and extracellular recordings from five nerves – *lvn, mvn, dgn, lgn*, and *ion* – are displayed. Horizontal lines indicate a reference membrane potential of -40 mV; vertical lines indicate a scale of 10 mV. Gray boxes correspond to the one-minute snapshots enlarged in subsequent panels respective to time. **(B-E)** Titles describe the pyloric neuron activity during and between LG bursts.

Over the 20 minutes shown, there was a clear increase in both gastric and pyloric activity with the LP and PD neurons becoming rhythmic and the emergence of strong rhythmic activity of the MCN1, LG, DG, and GM neurons (Figure 5A). At the beginning of this recovery period, both the PD and LP neurons were silent, reflecting loss of activity in pH 5.5 (Figure 5B). Strong LG neuron bursts were timed with hyperpolarizations of the LP neuron. This is consistent with previous findings that the neurons driving the gastric mill rhythm synapse onto the pyloric network and that gastric mill activity correlates with slowing of the pyloric rhythm (Bucher et al., 2007). A few minutes later, the LP and PD neurons started to recover rhythmic slow waves (Figure 5C). The LP and the second PD neuron – seen here on the *lvn* recording – were bursting. Both neurons became silent due to a strong inhibitory input coinciding with strong LG and MCN1 activity. Shortly thereafter, the LP and PD neurons were firing rhythmically (Figure 5D). Depolarizing inhibition resulted in tonic firing of LP and no activity on the intracellular recording of PD. Finally, the LP and PD neurons were bursting (Figure 5E). Inhibitory input coincident with LG and MCN1 activity resulted in depolarization of the PD and LP neurons and an increased duty cycle of LP bursting.

The rhythmic gastric-like activity seen here is similar to gastric mill rhythms elicited upon stimulation of the Ventral Cardiac Neurons (VCNs) (Beenhakker et al., 2007; Saideman et al., 2007; White and Nusbaum, 2011). Studies have shown that stimulation of the VCNs triggers activation of MCN1 and Commissural Projection Neuron 2 (CPN2) in the commissural ganglia (CoGs). This MCN1/CPN2 gastric mill rhythm drives the alternation of activity of the protractor motor neurons – LG, GM, MG, and IC – and the retractor neurons – DG, Int 1, and VD. We see similar activity here with strong MCN1 bursts on the *ion* corresponding to strong LG and GM bursts on the *lgn* and *dgn*, respectively, in alternation with DG bursts on the *dgn*.

### The cardiac rhythm is robust to acute changes in pH

To characterize the response of the cardiac rhythm to pH, we bath superfused cardiac ganglion preparations with saline between pH 5.5 and pH 10.4 using the same protocol described above for stomatogastric ganglion preparations. Example extracellular recordings are shown from the cardiac ganglia of two animals during the last minute of each pH step (Figure 6A). As shown in the top row of CG #1, the ganglion started in control saline at pH 7.8 and demonstrated a normal rhythm of Small and Large Cells bursting together. As the preparation was subjected to both acidic and basic saline, the rhythm remained. In contrast, the cardiac rhythm in CG #2 became less rhythmic in pH 5.5 and in pH 9.8 and above. A normal bursting rhythm recovered after superfusion of control saline as seen in the bottom right.

**Figure 6.**
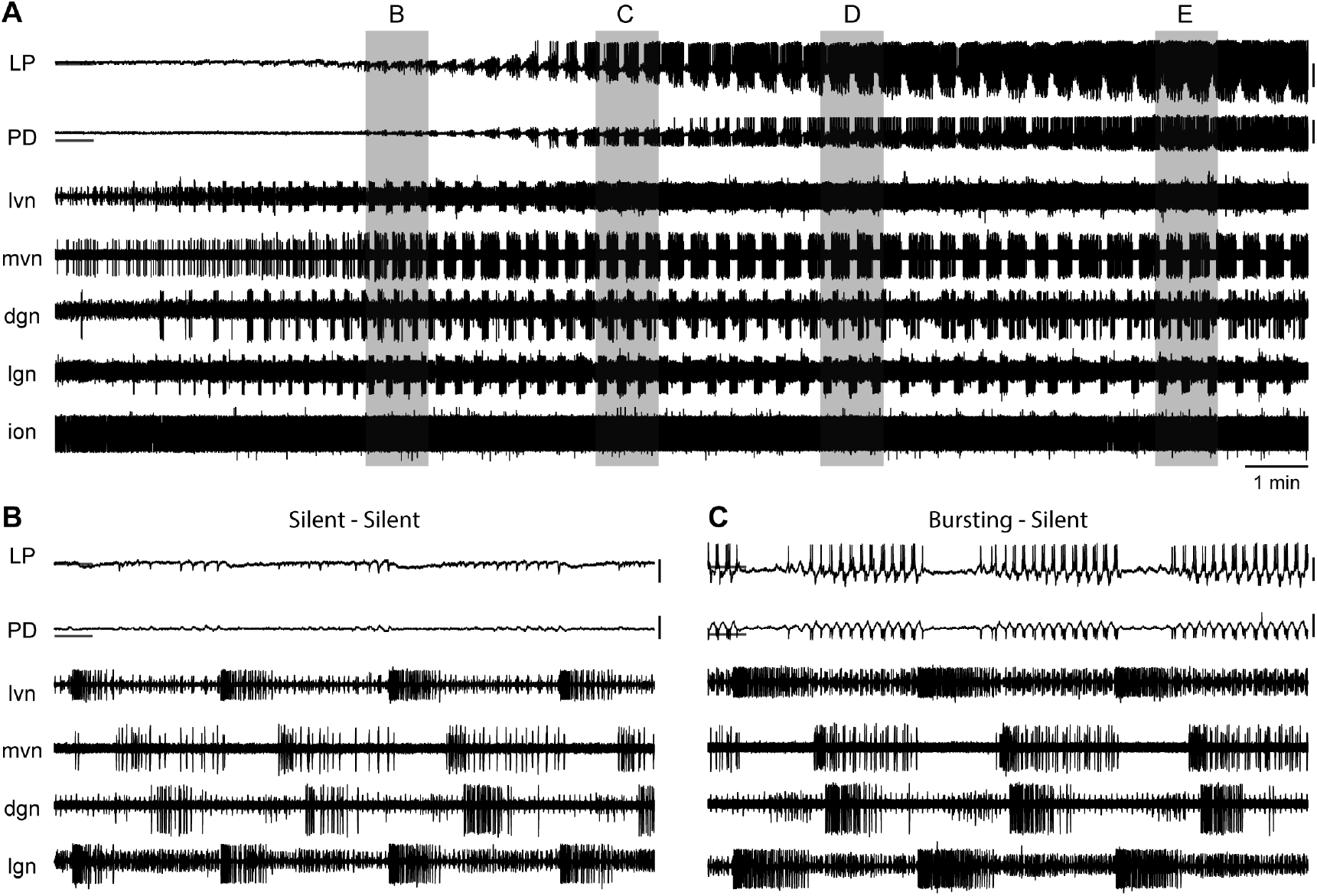
Robust and variable cardiac rhythm activity across pH. **(A)** Two example cardiac ganglion experiments with an acid-first protocol. Each colored box displays 12 seconds of extracellular recordings of the trunk taken from the last minute of each pH condition. Small Cell (SC) and Large Cell (LC) activity is visible. Each experiment can be read left to right then top to bottom in chronological order. **(B)** Cardiac frequency, **(C)** number of LC spikes per burst, and **(D)** LC duty cycle were calculated for CG #1 for each pH step. Violin plots show the KDE distribution, mean, median, IQR, and 95% CI for each measure across pH conditions. Recoveries from acid and base are displayed in the shaded gray regions on the far ends of each plot.

Measures of cardiac ganglion rhythm frequency, LC spikes per burst, and LC duty cycle were calculated for CG #1. Violin plots reveal the distribution of these measures at each pH (Figure 6B-D). The cardiac frequency of CG #1 decreased in acid and increased in base (Figure 6B). Further, the number of LC spikes per burst increased in acid (Figure 6C) while the LC duty cycle for CG #1 decreased slightly in acid and base (Figure 6D). Similar to STG #1, CG #1 retained robust activity throughout the entire range of pH tested.

To characterize these effects, we defined four states of activity: 1) ‘SC and LC bursting’ rhythm containing both units with a minimum of one LC spike per SC burst; 2) ‘SC bursting only’ rhythm containing only SC bursts with no or inconsistent LC spiking; 3) ‘all silent’; and 4) ‘atypical activity’ that could not be categorized under the first three definitions (Figure 7A). The mean fraction of time that all preparations spent in these states is plotted (Figure 7B). The mean fraction of time that preparations were rhythmic (state 1 – SC and LC bursting) was significant affected by both acid and base (Figure 7– figure supplement 1). Rhythmic activity was significantly decreased at pH 5.5, pH 9.3, pH 9.8, and pH 10.4 compared to control pH 7.8.

**Figure 7.**
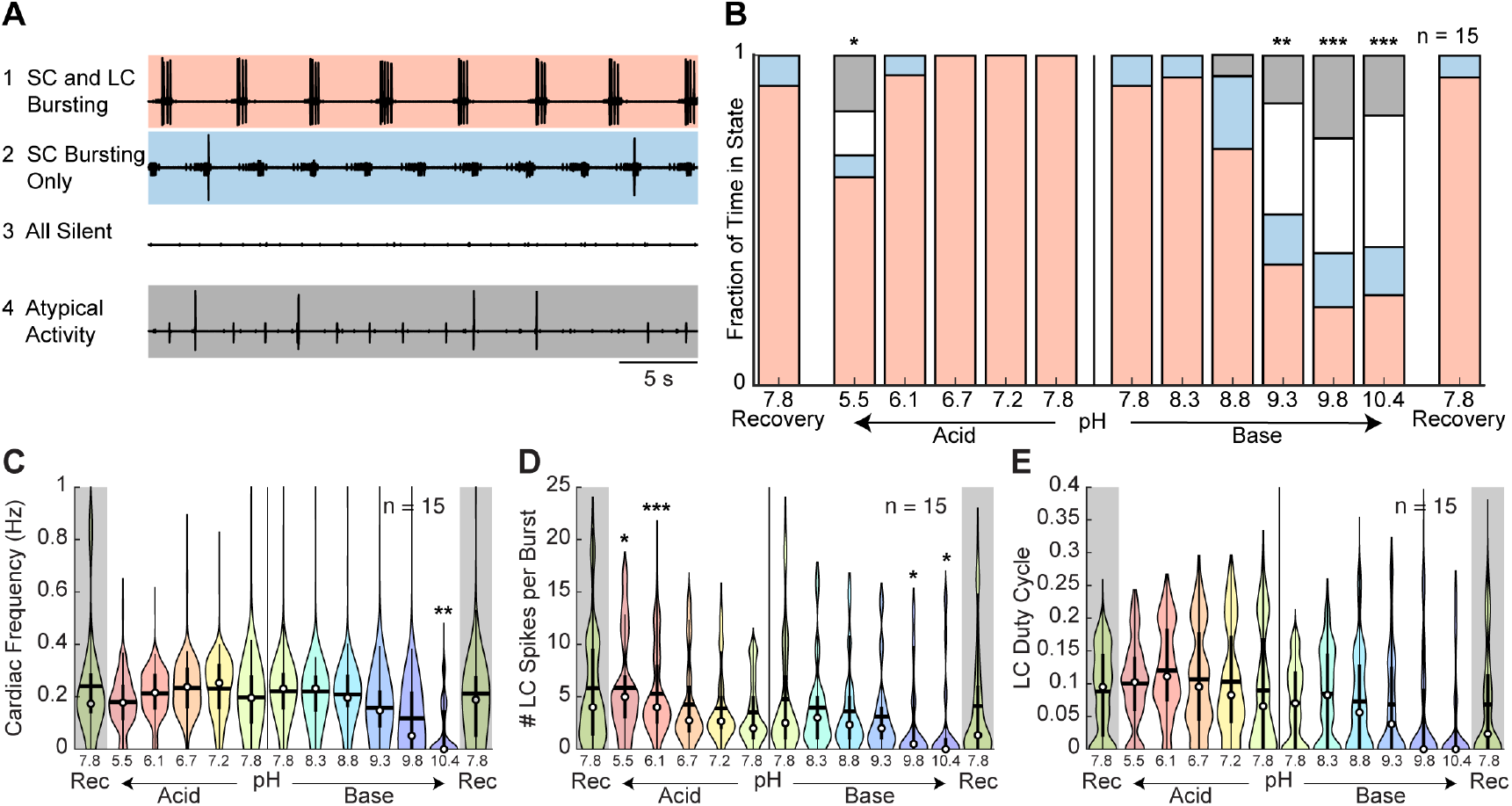
Characteristics of cardiac rhythm activity across pH. **(A)** Four states were defined to characterize cardiac rhythm activity. Examples of activity for each state are given. **(B)** Stacked bars give the mean fraction of time that all 15 preparations spent in each state for each pH step. **(C)** Cardiac rhythm frequency, **(D)** number of LC spikes per burst, and **(E)** LC duty cycle were calculated and pooled across all CG preparations for each pH step. Violin plots show the KDE distribution, mean, median, IQR, and 95% CI for each measure across pH conditions. Recoveries from acid and base are displayed in the shaded gray regions on the far ends of each plot. Asterisks denote statistical significance revealed by paired samples t-tests with Bonferroni correction (*p<0.05; **p<0.01; ***p<0.001).

To describe these effects quantitatively, measures of rhythm frequency, the number of LC spikes per burst, and LC duty cycle were calculated and their distributions are displayed in violin plots (Figure 7C-E). Cardiac rhythm frequency declined in both acidic and basic saline (Figure 7C; Figure 7–figure supplement 1). At pH 10.4, the cardiac rhythm was significantly slower. The mean number of LC spikes per burst was significantly affected in both acid and base (Figure 7D). The number of LC spikes per burst was significantly increased at pH 5.5 and pH 6.1 and decreased at pH 9.8 and pH 10.4. There was a significant effect of base but not acid on mean LC duty cycle (Figure 7E).

Similar to the STG, we observed a large spread in pooled measures across all pH conditions, reflecting the animal-to-animal variability in these networks. We plotted the distributions for all CG preparations at baseline and noted highly variable cardiac rhythm activity independent of the pH perturbation (Figure 7–figure supplement 2).

### The cardiac and pyloric rhythms are differentially sensitive to pH

To compare the effect of pH on the cardiac and pyloric rhythms, the distributions of the fraction of time that each preparation retained a normal rhythm were compared (Figure 8A). A normal rhythm was defined as a triphasic rhythm (a combined metric of states 1 and 2) and Small Cells and Large Cells bursting together (state 1) for the pyloric and cardiac rhythms, respectively. A comparison between the rhythmicity of these two ganglia across pH reveals similar distributions with maxima around control pH 7.8 and minima at extreme pH values. Interestingly, these distributions are asymmetrical, as the CG was more sensitive to extreme base whereas the STG was more sensitive to extreme acid. There were significant main effects of pH and ganglion as well as an interaction between pH and ganglion on rhythmicity in both acidic and basic solutions (Figure 8– figure supplement 1). The pyloric rhythm was significantly less rhythmic at pH 5.5, but significantly more rhythmic at 9.3 and 9.8 compared to the cardiac rhythm.

**Figure 8.**
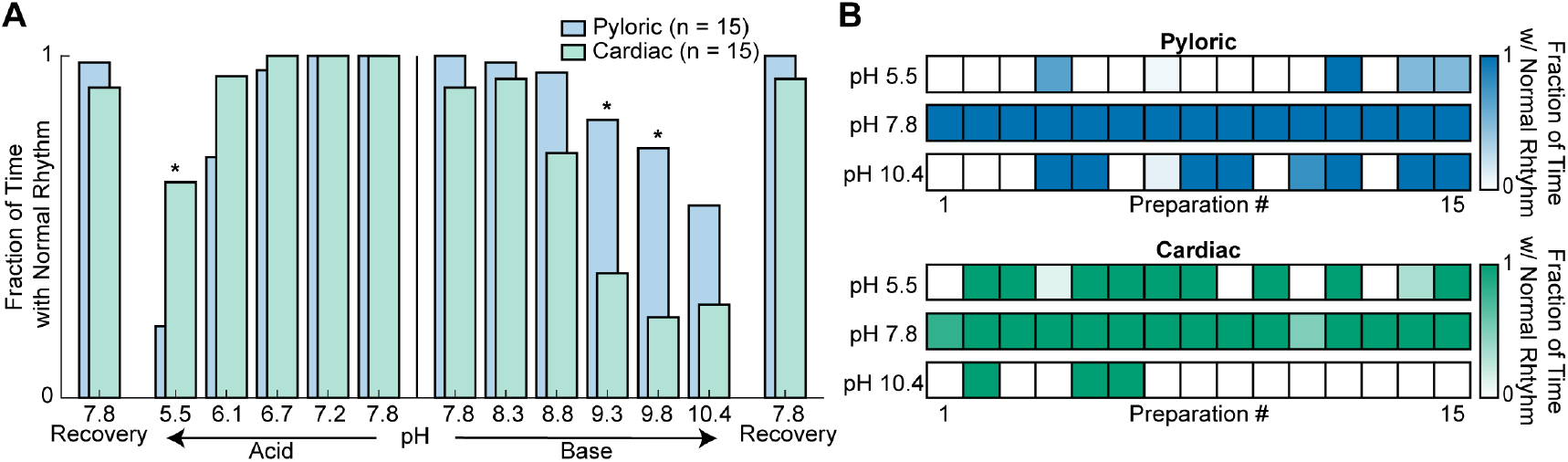
Rhythmicity of the cardiac and pyloric rhythms compared across pH. **(A)** Mean fraction of time that both the pyloric (blue) and cardiac (green) rhythms displayed normal activity is plotted as a function of pH. Differences between the activity of the two rhythms were analyzed by independent samples t-tests at each pH. Recovery from acid and base are displayed on the far ends of the plot. Asterisks denote statistical significance with Bonferroni correction (*p<0.05; **p<0.01; ***p<0.001). **(B)** Rhythmicity of individual animal preparations is plotted for extreme acid (pH 5.5), control (pH 7.8) and extreme base (pH 10.4) saline conditions. Each column of boxes represents a single preparation, with position across conditions remaining constant. The saturation of each box represents the mean fraction of time with a normal rhythm as indicated by the color bars on the right.

To understand better the amount of animal-to-animal variability in these two rhythms, the activity of individual preparations was plotted in control pH 7.8, extreme acid pH 5.5, and extreme base pH 10.4 (Figure 8B). Each box represents an individual preparation and its color saturation corresponds to the fraction of time with a normal rhythm. All preparations were rhythmically bursting at control pH – indicated by darkly colored boxes – and became less rhythmic – lighter colored – in the presence of extreme acid. However, 14 of 15 STG preparations ceased firing after 15 minutes of exposure to pH 5.5. In contrast, only 6 of 15 CG preparations showed reductions in rhythmic activity at pH 5.5. Three of the 15 CG preparations maintained a normal rhythm in every pH condition. Interestingly, STG preparations that showed decrements in activity during basic conditions were extremely susceptible to reductions in activity during extreme acid. The opposite is true in CG preparations suggesting that activity in base is a better predictor of acid activity in the STG and vice versa in the CG. This finding also suggests that some preparations were more susceptible to the effects of pH than others.

## Discussion

Circuit dynamics depend on the properties of the constituent neurons and their synaptic connections. Likewise, the intrinsic excitability of an individual neuron depends on the number and properties of its voltage- and time-dependent channels. Given that the physiological properties of ion channels are sensitive to pH (Anwar et al., 2017; Bayliss et al., 2015; Catterall, 2000; Cens et al., 2011; Cook et al., 1984; Doering and McRory, 2007; Guarina et al., 2017; Harms et al., 2017; Hille, 1968; Mahapatra et al., 2011; Marcanoti et al., 2010; Tombaugh and Somjen, 1996; Vilin et al., 2012; Zhou et al., 2018), one might imagine that a neuronal circuit might be as sensitive to changes in pH as its most sensitive ion channels. Therefore, it is surprising that both the pyloric rhythm of the STG and the cardiac ganglion rhythm in the crab, *C. borealis*, are relatively insensitive to acute pH change from about pH 6.1 to pH 8.8 while the individual functions of many ion channels are known to be considerably altered within this pH range.

One possibility is that crustacean ion channels are more robust to pH change than those that have been commonly studied. In most vertebrate animals, pH is carefully regulated. Slight acidosis or alkalosis can have deleterious effects on many aspects of vertebrate physiology (Chesler, 2003), which may be partially a consequence of the relative sensitivity of many vertebrate ion channels and synapses to pH. Unfortunately, little is known about the pH sensitivity of crustacean ion channels, but it would be surprising if it differed drastically from that seen in other animals as there is considerable homology across phylogeny in channel structure and function. However, it remains possible that modest evolutionary changes in channel structure occurred to allow endothermic animals to function in high temperature and low pH conditions.

A possible explanation for this circuit robustness may arise if there are compensatory and/or correlated changes in the effects of pH across the population of channels in these networks. Therefore, one prediction of the relative pH insensitivity of these networks is that numerous pH sensitive changes occur across the population of ion channels, but that these circuits have evolved sets of correlated ion channels that compensate for these changes (O’Leary and Marder, 2016; O’Leary et al., 2013; O’Leary et al., 2014; Temporal et al., 2012; Temporal et al., 2014; Tobin et al., 2009).

In addition to the relative pH insensitivity of these circuits, we were surprised that the cardiac and pyloric rhythms of *C. borealis* are differentially sensitive to acid and base. This was unexpected as many of the same ion channels are found in the two ganglia (Northcutt et al., 2016; Ransdell et al., 2013a; Ransdell et al., 2013b; Schulz et al., 2006; Schulz et al., 2007; Tobin et al., 2009). One possible explanation of this intriguing finding could be due to differences in the burst generating mechanisms of the two networks. The pyloric rhythm depends heavily on post-inhibitory rebound as a timing mechanism (Harris-Warrick et al., 1995a; Harris-Warrick et al., 1995b; Hartline and Gassie, 1979) while the cardiac ganglion depends on strong excitatory drive from the pacemaker neurons (Cooke, 1988). These excitatory and inhibitory synaptic connections could be differentially sensitive to pH. Additionally, although both networks are driven by bursting pacemaker neurons, the contribution of different ion channels to the burst generating mechanism may be sufficiently different in the two cases such that the pacemakers themselves respond differently to high and low pH.

We also found that the membrane potential of the isolated pyloric neurons, LP and PD, varied differentially with changing extracellular pH. Isolated LP neurons fired tonically and hyperpolarized in extreme base while isolated PD neurons depolarized in acid. In intact preparations, we observed depolarization in acid for both neurons, suggesting an important role of synaptic input in regulating network activity across pH. Together, these results illustrate the potential difficulty in generalizing the effects of environmental perturbation across neurons and circuits, even within the same animal.

Additionally, we found that under most control conditions, the gastric mill rhythm was silent as is typically observed in STG preparations. Unexpectedly, gastric mill rhythms were frequently activated upon exposure to or recovery from extreme pH. It is possible that either sensory or modulatory axons were activated by the pH changes, and it is feasible that sensory and/or modulatory neurons might be part of a response to altered pH.

In this study, we examined the effects of manipulating extracellular pH. However, the extent to which intracellular pH was affected and its contribution to changes in activity remain unclear. We observed that neurons penetrated with intracellular recording electrodes exhibited more labile activity in response to changing pH. This may indicate that changes in intracellular pH would be more deleterious than what occurs in response to changes in extracellular pH alone. Golowasch and Deitmer (1993) revealed that extracellular pH in the STG of the crab, *Cancer pagurus*, was reliably around 0.1 pH more alkaline than bath pH while intracellular pH was 0.3 to 0.4 pH more acidic. Further, moderate shifts in bath pH – from pH 7.4 to 7.0 or 7.8 – resulted in negligible changes in pyloric frequency and slow and low amplitude shifts in extracellular pH while NH_4_Cl induced acidosis resulted in recoverable alkylosis of both the intracellular and extracellular space (Golowasch and Deitmer, 1993). These results suggest the restriction of the free diffusion of protons through the ganglion and the existence of active Na^+^-dependent mechanisms to maintain more acidic intracellular and more alkaline extracellular compartments. Golowasch and Deitmer (1993) hypothesize that glial cells surrounding the neuronal processes in the neuropil of the STG may contain a Na^+^/H^+^ exchanger.

The ocean environment is both warming and acidifying at historic rates. *Cancer borealis* maintains relatively robust pyloric and cardiac rhythms in the temperature ranges it usually experiences, from 3°C to 25° C (Marder et al., 2015; Soofi et al., 2014; Tang et al., 2010; Tang et al., 2012). Importantly, the effect of temperature on ocean pH is relatively modest in comparison to the range of pH studied here. In *Carcinus meanus*, exposure to artificial ocean acidification produced relatively small changes in hemolymph pH (Maus et al., 2018). Therefore, unlike some ocean organisms that are very sensitive to even small ocean pH changes, we predict that the neuronal circuits in *C. borealis*, at least as an adult, will be largely insensitive to changes in ocean pH. However, other physiological parameters, such as metabolic rates and hemolymph flow may be more pH sensitive (Maus et al., 2018). Moreover, network performance may be further attenuated when pH is coupled to increasing temperature and other environmental insults.

Crab central pattern generating circuits are robust and adaptable to a large range of temperatures (2009; Haddad and Marder, 2018; Rinberg et al., 2013; Tang et al., 2010; Tang et al., 2012). Previous research revealed robust activity and increasing frequency of the pyloric rhythm of the STG and cardiac rhythm of the CG in response to increasing temperature in both *in vivo* and *ex vivo* preparations (Kushinsky et al., 2018; Tang et al., 2010; Tang et al., 2012). Contrastingly, increasing pH reveals non-linear effects on activity, which may suggest more complex mechanisms.

Finally, the data in this paper and in previous work on temperature reveal considerable animal-to-animal variability in response to extreme perturbations. Here, all preparations behaved predictably and reliably across more than a thousand-fold change in hydrogen ion concentration, an unexpectedly large range of robust performance. At more extreme pH, animal-to-animal variably became apparent, consistent with the responses of these circuits to extreme temperatures (Marder et al., 2015; Soofi et al., 2014; Tang et al., 2010; Tang et al., 2012). This animal-to-animal variability is almost certainly a consequence of the fact that similar network performance can arise from quite variable sets of underlying conductances (Goaillard et al., 2009; Grashow et al., 2009, 2010; Prinz et al., 2004). What remains to be seen is whether animals that are more robust to a given extreme perturbation are less robust to others and whether there are given sets of network parameters that confer robustness to many different perturbations.

## Materials and Methods

### Animals

From March 2016 to May 2018, adult male Jonah crabs (*Cancer borealis)* weighing between 400 and 700 grams were obtained from Commercial Lobster (Boston, MA). Before experimentation, all animals were housed in tanks with flowing artificial seawater (Instant Ocean) between 10°C and 13°C on a 12-hour light/dark cycle without food. Animals were kept in tanks for a maximum of two weeks. Animals were removed from tanks and kept on ice for 30 minutes prior to dissection.

### Saline Solutions

Control *C. borealis* physiological saline was composed of 440 mM NaCl, 11 mM KCl, 13 mM CaCl_2_, 26 mM MgCl_2_, 11 mM Trizma base, and 5 mM Maleic acid. Additional quantities of concentrated HCl and NaOH were added to achieve solutions with pH 5.5, 6.1, 6.7, 7.2, 7.8, 8.3, 8.8, 9.3, 9.8, and 10.4, at 11°C. Solution pH was measured using a calibrated pH/ion meter (Mettler Toledo S220). For experiments with picrotoxin, 10^-5^ M PTX was added to each of the pH solutions.

### Electrophysiology

The stomatogastric and cardiac nervous systems were dissected out of the animals and pinned out in a Sylgard (Dow Corning) coated plastic Petri dish containing chilled saline (11°C). In all cases, we worked only with fully intact stomatogastric nervous system preparations that included the commissural and esophageal ganglia and their descending nerves. Only preparations containing healthy cardiac or pyloric rhythms with no sign of damage from dissection were analyzed.

Vaseline wells were placed around motor nerves and extracellular recordings were obtained using stainless steel pin electrodes placed in the wells and amplified using a differential amplifier (A-M Systems Model 1700). Intracellular sharp-electrode recordings were obtained from cell bodies in the stomatogastric ganglion using a microelectrode amplifier (Molecular Devices Axoclamp 2B or 900A) with HS-2A-x1LU headstages holding 15-30 MΩ boroscilate microelectrodes with filaments (Sutter Instrument Co. BF150-86-10) pulled with a Flaming/Brown micropipette puller (Sutter Instrument Co. P-97). Microelectrodes were filled with a solution of 10 mM MgCl_2_, 400 mM potassium gluconate, 10 mM HEPES, 15 mM Na_2_SO_4_, and 20 mM NaCl (Hooper et al., 2015).

Preparations were continuously superfused with physiological saline at 11°C. Superfusion was gravity fed at approximately nine mL/min. The temperature of the superfusing saline was controlled and recorded using a Peltier device (Warner Instruments CL-100). Instantaneous bath pH was recorded using a pH microelectrode placed adjacent to the ganglion (Thermo Scientific Orion 9810BN) combined with a preamplifier (Omega PHTX-21). Output from the pH microelectrode was converted from arbitrary voltage to pH using a temperature-compensated calibration.

### Data Acquisition and Analysis

Data were acquired using a data acquisition board (Molecular Devices Digidata 1440A) and Clampex 10.5 software (Molecular Devices). Data were analyzed using Clampfit 10.5, Spike2 v 6.04 (Cambridge Electronic Design), and/or MATLAB 2017A (MathWorks). All code is available for download at github.com/jesshaley/haley_2018. Figures were prepared in Adobe Illustrator CC 2017.

For analyses of extracellular recordings of the *lvn* of the STG or the trunk of the CG, we analyzed the last eight minutes of each 15-minute pH step to ensure that preparations had reached a steady state.

Data were categorized into states by manual annotation. A transition from one state to another was noted when there was a sustained change in activity lasting a minimum of 10 seconds. In other words, if the rhythm transitioned from one state into another and maintained the new state of activity for at least 10 seconds, a transition was noted at the start of that new state. Rhythms rarely transitioned intermittently between two states (e.g. once a pyloric rhythm had transitioned from normal triphasic to weak triphasic, it rarely transitioned back to normal triphasic until after recovery in control pH). Further, rhythms generally transitioned in a stereotypical pattern. For STG preparations, the pyloric rhythm often transitioned from normal triphasic to weak triphasic to intermittent triphasic to all silent. For CG preparations, the cardiac rhythm usually transitioned from SC and LC bursting to SC bursting only to all silent. During recovery, these transition patterns were reversed. The mean fraction of time that the preparations remained in each state during the last eight minutes of recording at each pH step is plotted as stacked bar graphs.

Quantitative variables of frequency, number of spikes per burst, and duty cycle were measured using extracellular recordings. Spikes and bursts were first isolated in Spike2 by thresholding extracellular recordings. MATLAB was then used for further analysis. Instantaneous burst frequency was calculated by taking the inverse of the cycle’s period, the time elapsed between the onset of one burst and the onset of the next. The number of spikes per burst of a given neuron reflects the number of spikes contributing to each burst. Duty cycle reflects the fraction derived by dividing the burst duration – time elapsed between the first and last spike – by the burst period. Mean values were computed for bins of 10 seconds such that for eight minutes of data, there were 48 binned mean values for each preparation, condition, and measure. Violin plots show distributions of these binned mean values pooled for all preparations. The body of the violin is a rotated kernel density estimate (KDE) plot. The circles give the median of the pooled data and the horizontal bars give the mean. The interquartile range (IQR) is given by the box plot within each violin with the whiskers giving the 95 percent confidence interval (CI).

For analyses of intracellular recordings of isolated LP and PD neurons, the last minute of each pH step was analyzed in MATLAB. Minimum membrane potential was first measured by finding the minimum voltage of the neuron between each burst. Recordings were then low-pass filtered to remove spikes from the slow wave. Slow wave amplitude was measured by subtracting the trough from the peak of the slow wave’s membrane potential. Spike amplitude was retrieved by subtracting the filtered slow wave signal from the original recording and then measuring the amplitude from trough to peak of each action potential. PD burst frequency was calculated by finding the inverse of the time period between one slow wave trough to the next. LP firing rate was determined by calculating the inverse of the inter-spike interval, the time between spikes. Mean values were computed for bins of 10 seconds. Violin plots show distributions of these binned mean values pooled for all preparations.

### Statistics

All statistics were performed using R (version 3.4.3). We performed statistical testing of the effects of acid and base on measures of the cardiac and pyloric rhythms using a Univariate Type III Repeated-Measures Analysis of Variance (ANOVA) from the car package. Separate tests were performed for acid and base step protocols. Post-hoc paired sample t-tests with Bonferroni correction were performed for each pH step against its respective control. To assess the differences between the effects of pH on the cardiac and pyloric rhythms, we performed a Two-Way Mixed-Measures ANOVA (Type III) for both acid and base step protocols using the car package. Post-hoc independent samples t-tests with Bonferroni correction were performed for each pH condition.

---

**Figure 3–figure supplement 1.**
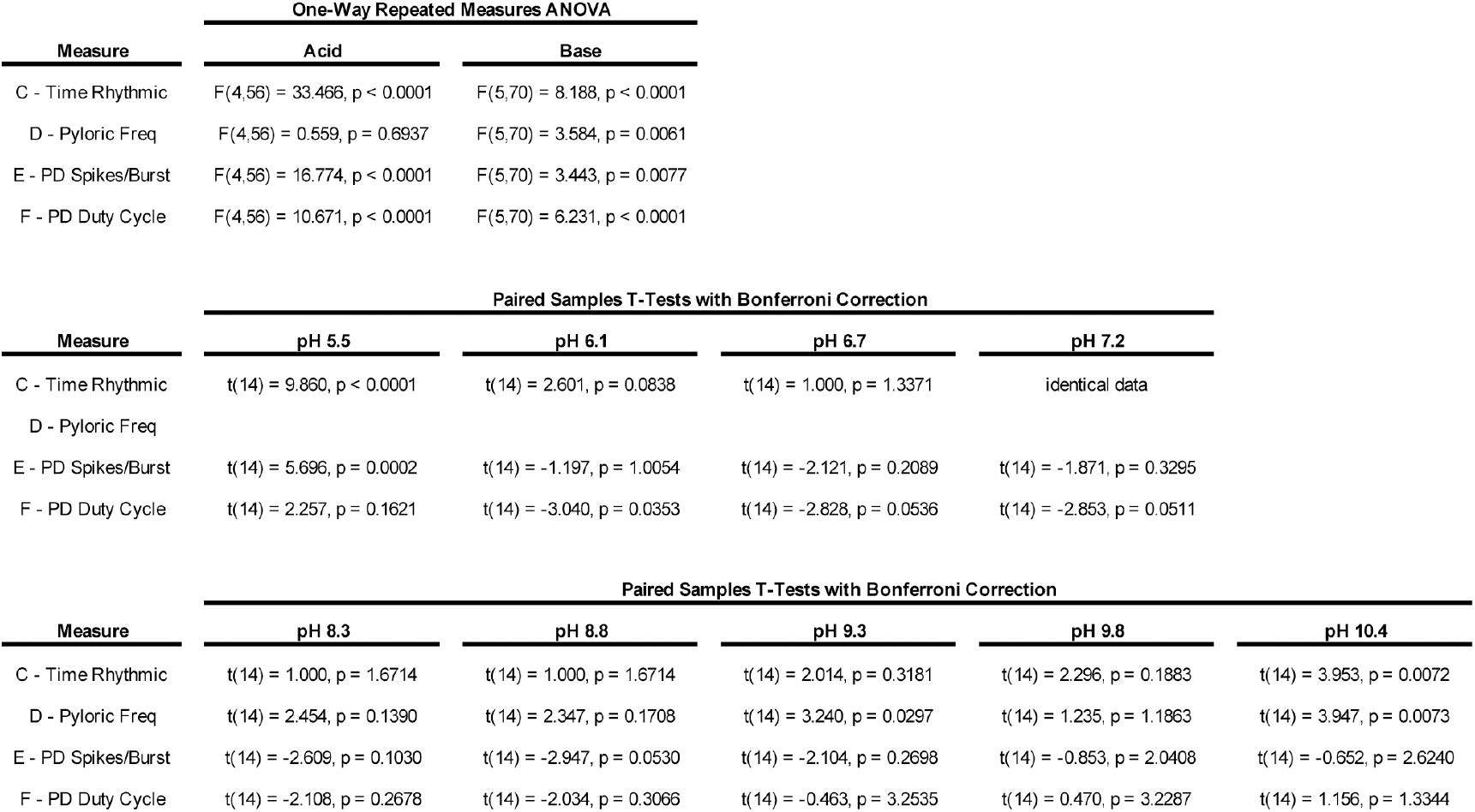
Statistical analysis of the effects of pH on the pyloric rhythm. The main effects of acid and base protocols on four measures of the activity of the pyloric rhythm were assessed. Univariate Type III Repeated-Measures Analysis of Variance (ANOVA) tests were performed separately for both acid and base step protocols. Post-hoc paired samples t-tests with Bonferroni correction were performed for each pH step against its respective control, the pH 7.8 condition immediately prior to the step protocol.

**Figure 3–figure supplement 2.**
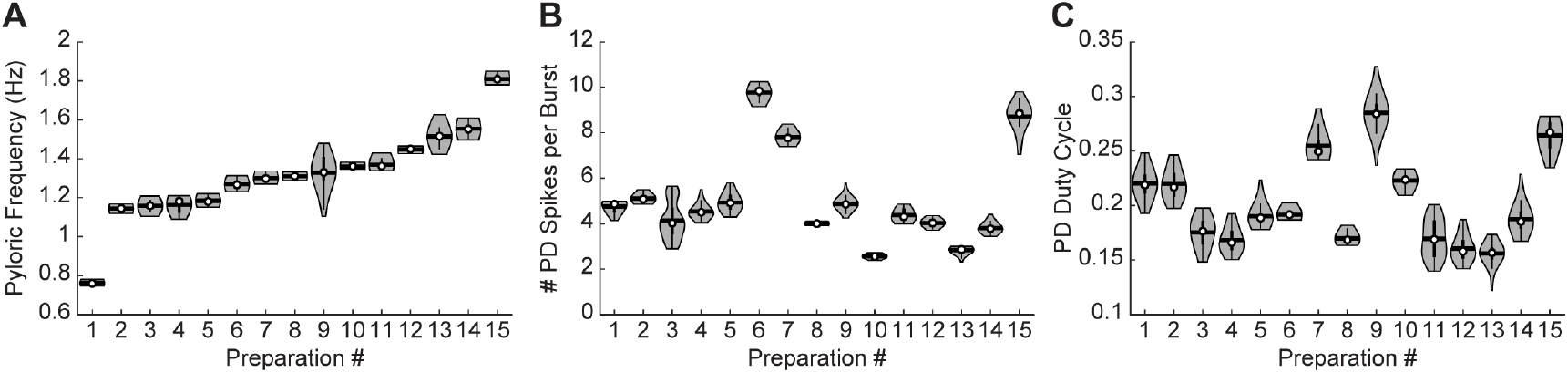
Variability of pyloric rhythm activity at control pH. **(A)** Pyloric rhythm frequency, **(B)** number of PD spikes per burst, and **(C)** PD duty cycle were calculated for each CG preparation for the last eight minutes in control pH 7.8. Violin plots show the KDE distribution, mean, median, IQR, and 95% CI for each measure across preparations. Preparations are consistent across plots and sorted in order of increasing frequency.

**Figure 4–figure supplement 1.**
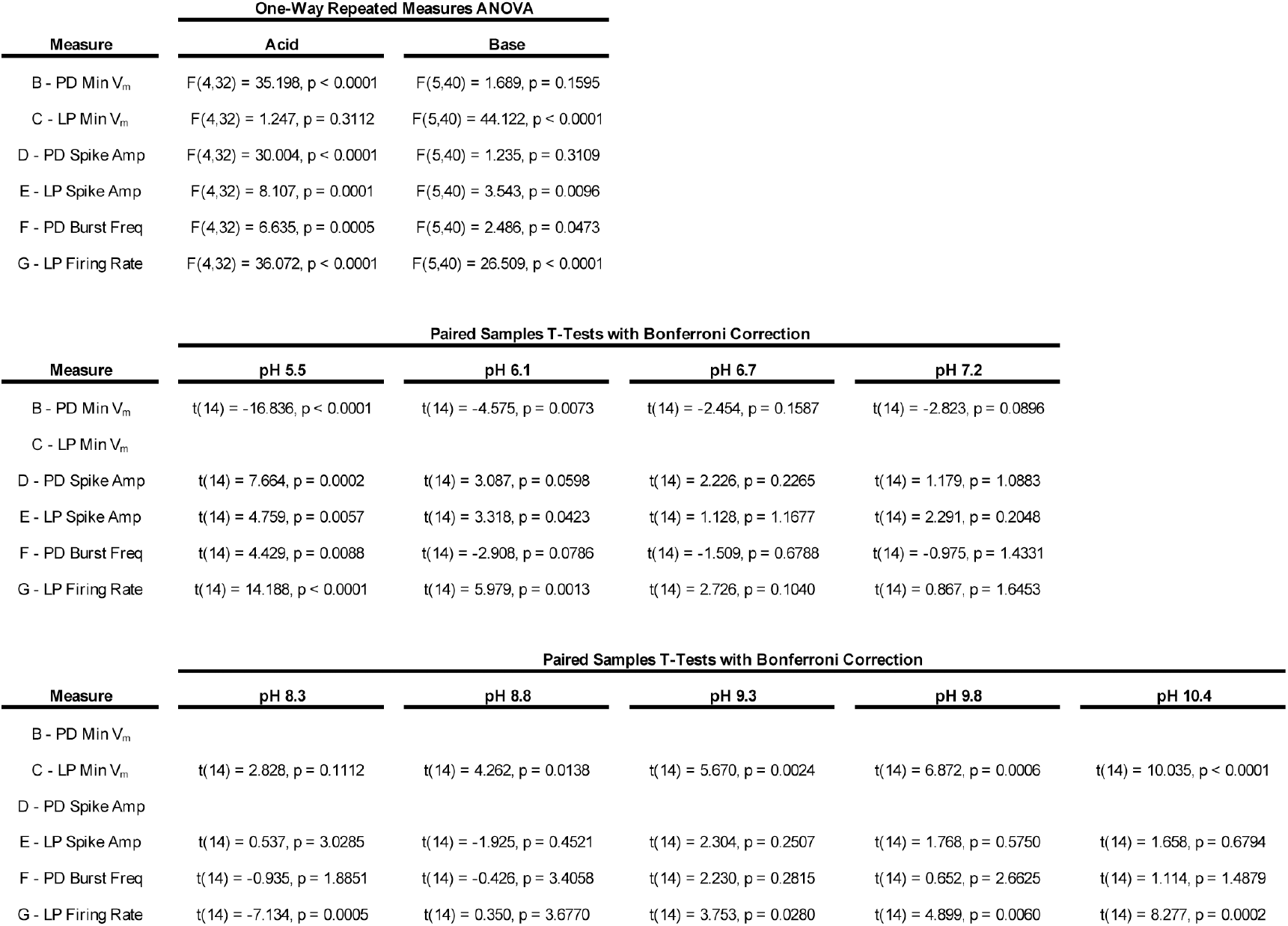
Statistical analysis of the effects of pH on isolated PD and LP neurons. The main effects of acid and base protocols on six measures of the activity of isolated LP and PD neurons were assessed. Univariate Type III Repeated-Measures Analysis of Variance (ANOVA) tests were performed separately for both acid and base step protocols. Post-hoc paired samples t-tests with Bonferroni correction were performed for each pH step against its respective control, the pH 7.8 condition immediately prior to the step protocol.

**Figure 7–figure supplement 1.**
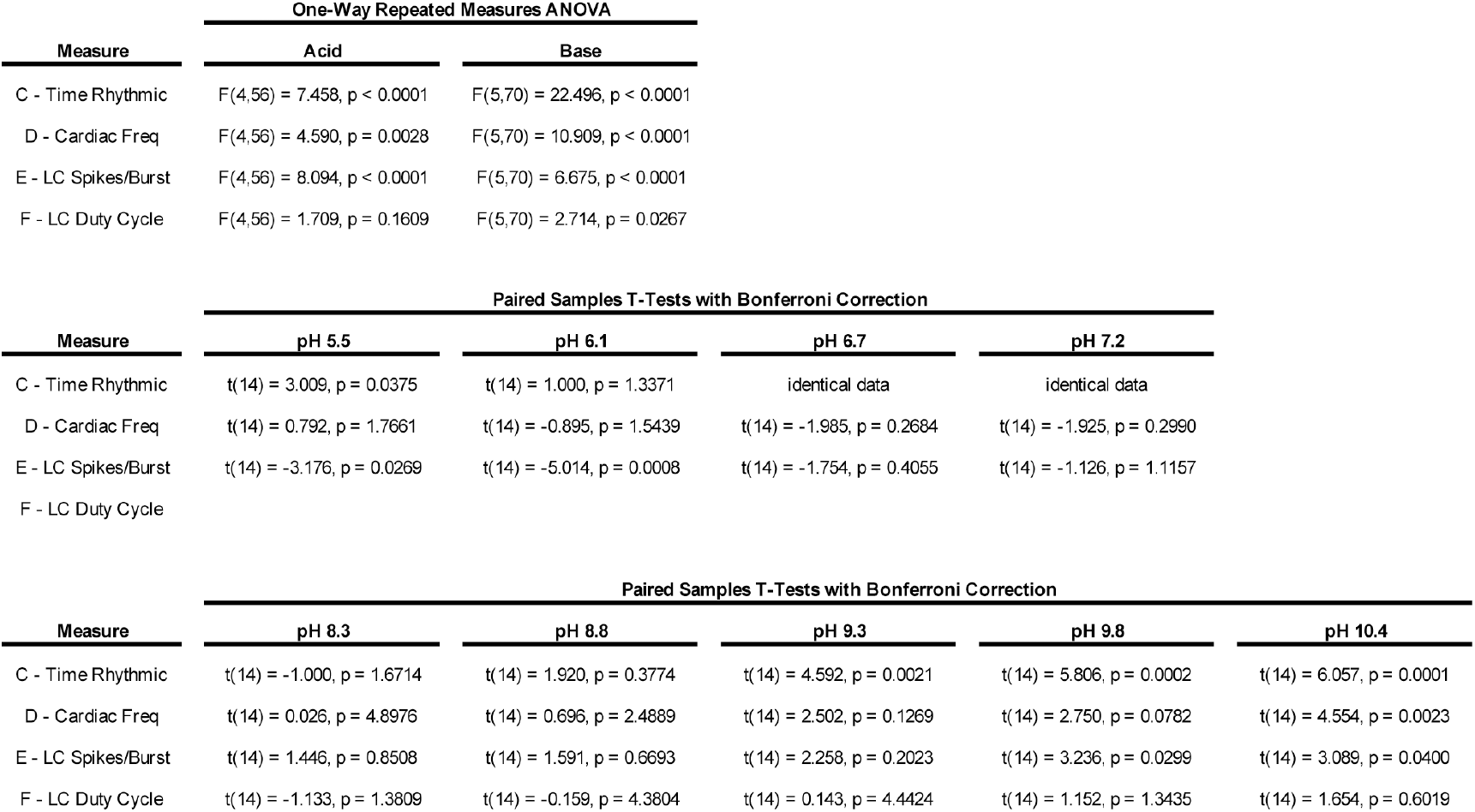
Statistical analysis of the effects of pH on the cardiac rhythm. The main effects of acid and base protocols on four measures of the activity of the cardiac rhythm were assessed. Univariate Type III Repeated-Measures Analysis of Variance (ANOVA) tests were performed separately for both acid and base step protocols. Post-hoc paired samples t-tests with Bonferroni correction were performed for each pH step against its respective control, the pH 7.8 condition immediately prior to the step protocol.

**Figure 7–figure supplement 2.**
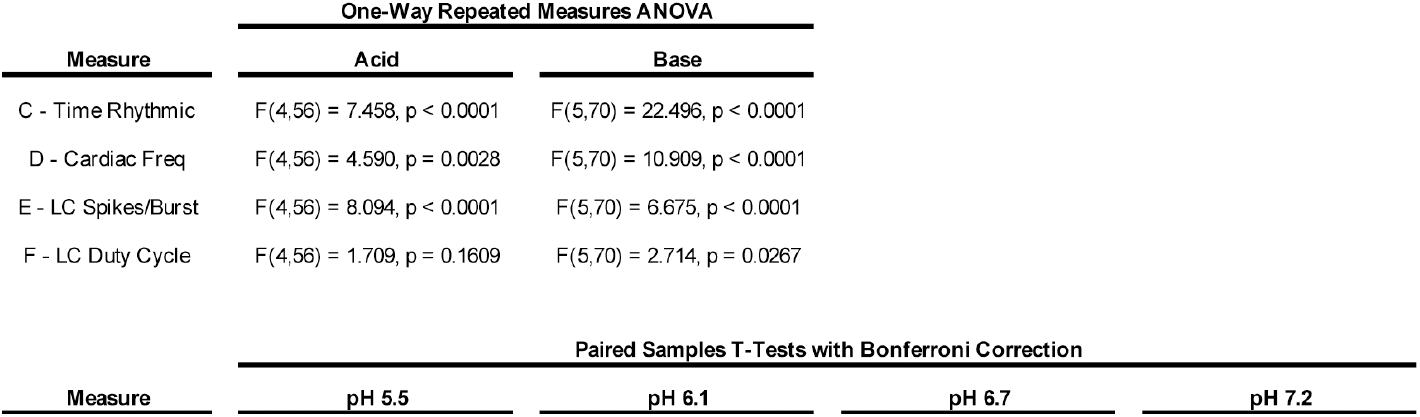
Variability of cardiac rhythm activity at control pH. **(A)** Cardiac rhythm frequency, **(B)** number of LC spikes per burst, and **(C)** LC duty cycle were calculated for each CG preparation for the last eight minutes in control pH 7.8. Violin plots show the KDE distribution, mean, median, IQR, and 95% CI for each measure across preparations. Preparations are consistent across plots and sorted in order of increasing frequency.

**Figure 8–figure supplement 1.**
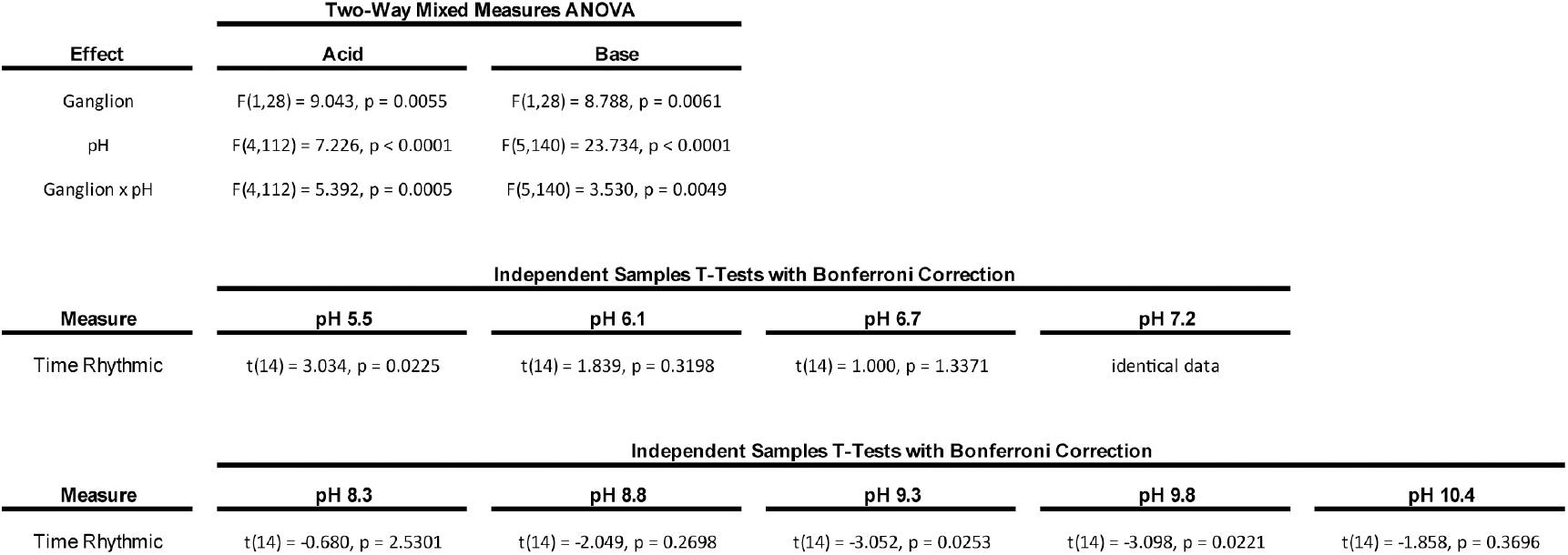
Statistical analysis of the differential effects of pH on the pyloric and cardiac rhythms. The main effects of ganglion and pH and their interaction during both acid and base protocols on the fraction of time rhythmic of both the pyloric and cardiac rhythms were assessed. Multivariate Type III Mixed-Measures Analysis of Variance (ANOVA) tests were performed separately for acid and base step protocols. Post-hoc independent samples t-tests with Bonferroni correction were performed to compare rhythmicity of stomatogastric and cardiac preparations at each pH step.

Author contributions
Jessica A. Haley, Data curation, Formal Analysis, Investigation, Methodology, Software, Validation, Visualization, Writing – Original Draft Preparation, Writing– Review & Editing; David Hampton, Formal Analysis,Investigation, Writing – Original Draft Preparation, Writing – Review & Editing; Eve Marder, Conceptualization, Data Curation, Funding Acquisition, Project Administration, Resources, Supervision, Writing – Review & Editing

